# Polygenic risk score for schizophrenia is more strongly associated with ancestry than with schizophrenia

**DOI:** 10.1101/287136

**Authors:** David Curtis

**Affiliations:** UCL Genetics Institute, UCL, Darwin Building, Gower Street, London WC1E 6BT; Centre for Psychiatry, Barts and the London School of Medicine and Dentistry, Charterhouse Square, London EC1M 6BQ

**Keywords:** Schizophrenia, polygenic risk score, RNA, expression

## Abstract

**Background:** The polygenic risk score (PRS) for schizophrenia, derived from very large numbers of weakly associated genetic markers, has been repeatedly shown to be robustly associated with schizophrenia in independent samples and also with other diseases and traits.

**Aims:** To explore the distribution of the schizophrenia PRS in subjects of different ancestry.

**Method:** The schizophrenia PRS derived from the large genome-wide association study carried out by the Psychiatric Genetics Consortium was calculated using the downloaded genotypes of HapMap subjects from eleven different ancestral groups. It was also calculated using downloaded genotypes of European schizophrenia cases and controls from the CommonMind Consortium.

**Results:** The PRS for schizophrenia varied significantly between ancestral groups (p < 2*10^−16^) and was much higher in African than European HapMap subjects. The mean difference between these groups was ten times as high as the mean difference between European schizophrenia cases and controls. The distributions of scores for African and European subjects hardly overlapped.

**Conclusions:** The PRS cannot be regarded as simply a measure of the polygenic contribution to risk of schizophrenia and clearly contains a strong ancestry component. It is possible that this could be controlled for to some extent by incorporating principal components as covariates but doubts remain as to how it should be interpreted. The PRS derived from European subjects cannot be applied to non-Europeans, limiting its potential usefulness and raising issues of inequity. Previous studies which have used the PRS should be re-examined in the light of these findings.

**Declaration of interest:** The author declares he has no conflict of interest.

## Introduction

The polygenic risk score (PRS) is derived by estimating the effect size of large numbers of genetic variants from a case-control discovery sample and combining these across the genotypes observed in other subjects (Visscher *et al*., 2017). Typically, many thousands of variants are used to contribute to the PRS with the expectation that only a proportion of them will in fact be truly associated with the trait in question. The first major application of this approach was to a genome wide association study (GWAS) of schizophrenia and was used to justify the claim that thousands of alleles contribute to the risk of schizophrenia and that there is shared genetic risk between schizophrenia and bipolar disorder (Purcell *et al*., 2009). Since this landmark publication, the observation that the PRS derived from one sample is associated with schizophrenia in another sample has been replicated numerous times and is probably one of the most robust observations in psychiatric genetics. The effect size is substantial and in a larger GWAS the odds ratio for schizophrenia risk between subjects with PRS in the highest or lowest decile was around 10 (Schizophrenia Working Group of the Psychiatric Genomics Consortium, 2014). This study made summary statistics available for each marker and this would allow other researchers to use it as a discovery sample to calculate the PRS in other samples. However it should be noted that in order to carry out a GWAS cases and controls need to be matched for ancestry because marker allele frequencies vary between populations. Thus the summary statistics were produced from a meta-analysis of a number of matched case-control samples which were overwhelmingly of European ancestry. Of a total of 38,131 schizophrenia cases and 114,674 controls, all were of European ancestry except 1,866 cases and 3,418 controls who were of East Asian ancestry. Using the summary statistics and applying them to other traits allows one to test whether the variants associated with increased risk of schizophrenia are also associated with other disorders and a recent review identified 31 articles examining the association of the schizophrenia PRS with other psychiatric and non-psychiatric phenotypes (Mistry *et al*., 2017).

Given the way that it is derived and how it is used, there is a natural tendency to assume that the PRS produced for a trait is a measure of polygenic susceptibility to that trait. However a recent study of autism showed in a supplementary figure that the PRS for schizophrenia stratified by ancestry (Weiner *et al*., 2017) and in a separate study investigating whether the PRS predicted gene expression it was noted that the PRS for schizophrenia was strongly correlated with the first principal component of marker genotypes (a proxy for ancestry) in both cases and controls of the Common-Mind Consortium (CMC) dataset (Curtis, 2017). Similarly, a recent study demonstrated that the PRS for both type 2 diabetes and coronary heart disease varies between populations and pointed out that this would need to be taken into account if attempting to estimate an individual’s risk (Reisberg *et al*., 2017).

In order to investigate this more formally, the present study set out to examine the distribution of the PRS in cohorts of different ancestry genotyped for the HapMap project and to compare this with the distribution between schizophrenia cases and controls in the CMC dataset which had initially revealed the principal component correlation.

## Methods

The merged post-QC phase I+II and III HapMap (International HapMap 3 Consortium *et al*., 2010) genotype files were downloaded from ftp://ftp.ncbi.nlm.nih.gov/hapmap/genotypes/2010-08_phaseII+III/forward/. These subjects are from eleven different ancestral groups and are assumed not to be affected with schizophrenia. Since the prevalence of schizophrenia is only 1% it seems reasonable to assume, even in the absence of a formal psychiatric assessment, that at least the vast majority are unaffected. In order to obtain a PRS for schizophrenia, the file called *scz2.prs.txt.gz*, containing ORs and p values for 102,636 LD-independent single nucleotide polymorphism markers (SNPs), was downloaded from the Psychiatric Genetics Consortium (PGC) website (*www.med.unc.edu/pgc/results-and-downloads*). The ORs had been obtained by carrying out metanalysis of 49 cohorts of European ancestry along with 3 cohorts of East Asian ancestry and using principal components to control for population stratification. This training set was produced as part of the previously reported PGC2 schizophrenia GWAS (Schizophrenia Working Group of the Psychiatric Genomics Consortium, 2014). SNPs from this dataset were then selected only if they had also been genotyped in all 11 of the HapMap cohorts, yielding a reduced set of 32,588 SNPs. HapMap subjects with genotyping call rate < 0.90 were removed, leaving a sample of 1,397.

The dataset used in the previous gene expression studies was downloaded from the CMC Knowledge Portal (*https://www.synapse.org/#!Synapse:syn2759792/wiki/69613*) consisting of SNP genotypes and RNAseq results from frontal cortex samples originating from tissue collections at Mount Sinai NIH Brain Bank and Tissue Repository (MSSM), University of Pennsylvania Brain Bank of Psychiatric illnesses and Alzheimer’s Disease Core Center (Penn) and The University of Pittsburgh NIH NeuroBioBank Brain and Tissue Repository (Pitt), collectively referred to as the CMC MSSM-Pitt-Penn dataset (Fromer *et al*., 2016). Genotypes and expression levels were available for 258 subjects with schizophrenia and 279 controls, though in the current analysis only the genotypes were used. The distributions of ethnicities were reported to be similar between subjects with schizophrenia and controls (Caucasian 80.7%, African-American 14.7%, Hispanic 7.7%, East Asian 0.6%). The methods for obtaining the genotypes and expression data have been described by the authors of the original study (Fromer *et al*., 2016). Genotyping was performed on the Illumina Infinium HumanOmniExpressExome 8 v 1.1b chip (Catalog #: WG-351-2301) using the manufacturer’s protocol. QC was performed using PLINK to remove markers with: zero alternate alleles, genotyping call rate < 0.98, Hardy-Weinberg p-value < 5 × 10^−5^, and individuals with genotyping call rate < 0.90. Marker alleles were phased to the forward strand, and ambiguously stranded markers were removed.

SNPs with p value < 0.05 in the PGC2 training set were selected and their log(OR) summed over sample genotypes using the *--score* function of *plink 1.09beta* in order to produce a PRS for each subject in both the HapMap and CMC datasets (www.cog-genomics.org/plink/1.9/) (Purcell *et al*., 2007, 2009; Chang *et al*., 2015). The first 20 principal components for both datasets were produced using the *--pca* and *--make-rel* functions of *plink*.

As described previously (Curtis, 2017) ancestral outliers were removed from the CMC dataset by removing subjects with values for the first or second principal component exceeding −0.01. This left a fairly homogeneous sample of 264 subjects in which the PRS was not correlated at p < 0.05 with 19 of the first 20 principal components, although it was correlated with the 11th at p = 0.0019.

Statistical tests and data manipulation were carried out using R version 3.3.2 (R Core Team, 2014). The distributions of the PRS were compared between HapMap cohorts using anova and additionally a correlation analysis of PRS with the principal components was performed. For the CMC dataset, a t test was used to compare the PRS between subjects with schizophrenia and controls. In both samples, the residuals of the PRS after correction for the first 20 principal components were also compared.

## Results

The distribution of schizophrenia PRS between HapMap cohorts is shown in Table 1 and Figure 1. The SD of the PRS is fairly similar in all cohorts, varying from 1.71 to 2.59. However the means are very different. In CEU and TSI, the cohorts of European ancestry, the average PRS is −2.90 and −2.64. In ASW, LWK, MKK and YRI, the cohorts of African ancestry, the average PRS is 7.75, 9.48, 7.08 and 10.27. As can be seen from Figure 1, the differences between the cohorts are so marked that the scores for the European and African cohorts scarcely overlap. The anova testing for a difference between cohort means was formally statistically significant at p < 2*10^−16^. In the correlation analysis with the first 20 principal components, the PRS was highly significantly correlated with each of the first 6 principle components, at values of p < 2*10^−12^ or lower. The residual PRS from this analysis did not significantly differ between cohorts.

**Table 1.** Schizophrenia PRS distribution in HapMap cohorts.

| Cohort | Description | Number of subjects | PRS mean (SD) |
| --- | --- | --- | --- |
| ASW | African ancestry in Southwest USA | 87 | 7.75 (2.05) |
| CEU | Utah residents with Northern and Western European ancestry | 165 | -2.90 (2.09) |
| CHB | Han Chinese in Beijing, China | 137 | 4.71 (1.78) |
| CHD | Chinese in Metropolitan Denver, Colorado | 109 | 4.57 (1.71) |
| GIH | Gujarati Indians in Houston, Texas | 101 | 1.91 (2.08) |
| JPT | Japanese in Tokyo, Japan | 113 | 4.71 (1.59) |
| LWK | Luhya in Webuye, Kenya | 110 | 9.48 (1.76) |
| MEX | Mexican ancestry in Los Angeles, California | 86 | 1.50 (2.59) |
| MKK | Masai in Kinyawa, Kenya | 184 | 7.08 (1.76) |
| TSI | Toscani in Italy | 102 | -2.64 (1.91) |
| YRI | Yoruba in Ibadan, Nigeria | 203 | 10.27 (1.65) |

In the CMC dataset, the PRS was significantly higher in subjects with schizophrenia (mean – 5.14) than controls (mean −6.08), difference = 0.94, p = 2.5*10^−7^. The residual PRS after correcting for the first 20 principal components in this dataset remained significantly higher in the subjects with schizophrenia, p = 7.4*10^−7^, though with a somewhat smaller difference between the means (0.41 versus −0.47, difference = 0.88).

## Discussion

There are striking differences in the schizophrenia PRS between cohorts with different ancestries. The differences between subjects of European and African ancestry are much larger, by a factor of around 10, than the differences between subjects with schizophrenia and controls of European ancestry.

The underlying mechanisms responsible for producing these results are not immediately obvious. Although differences in the frequencies of individual variants between groups of different ancestries would be not unexpected, it is harder to see why there should be a systematic effect such that alleles found more commonly in European schizophrenic subjects in the PGC should generally tend to be commoner in subjects with African ancestry. Two kinds of explanation suggest themselves. The most benign, from the point of view of the usefulness of the PRS, is that the PRS does indeed indicate genetic susceptibility to schizophrenia and that the contributing alleles are under stronger negative selection in African than non-African environments. The least benign would be to say that the PRS is basically an indicator of African ancestry and that for some reason, perhaps through mechanisms such as social adversity, subjects in the PGC with schizophrenia have a higher African ancestry component than controls. It does not seem that the latter can be a full explanation, because it does seem that the PRS is associated with schizophrenia risk in a homogeneous sample after correction for principal components. On the other hand, it is difficult to accept that the PRS does not index ancestry to at least some extent. Although there is some evidence that rates of schizophrenia are higher among subjects with African ancestry resident, for example, in the UK (Pinto, Ashworth and Jones, 2008), the prevalence appears to be fairly similar across the world and if anything lower in developing countries (Bhugra, 2005). Thus it does not seem plausible that the genetic risk associated in subjects with African ancestry could be as high as a naïve interpretation of the PRS would imply. This finding also impacts on the observation that for complex traits very large numbers of association signals are detected across the genome – the so-called “omnigenic” effect (Boyle *et al*., 2017). This is what would be expected if complex trait associations were in fact signals of ancestry rather than of disease risk.

Whatever the explanation, these results have important implications for the interpretation of the PRS. As has been pointed out in the case of type 2 diabetes and coronary heart disease, the fact that the PRS varies according to ancestry would raise problems were one to attempt to use it to estimate an individual’s risk in a clinical situation (Reisberg *et al*., 2017). Any proposal to use the PRS to predict risk in individual subjects would need to explain very clearly how ancestry issues were to be dealt with. For certain applications, the PRS may be adjusted using principal components and other measures but given the very strong ancestry effects it would be reasonable to be concerned about whether such adjustments could ever be fully effective at the level of the individual. It may be worth noting that the PRS used for this study was obtained from analyses of mostly European samples in which principal components had already been incorporated but that nevertheless the resulting measure still correlated very strongly with the principal components of an ancestrally heterogeneous sample. It might be that the PRS is a measure of relative polygenic risk in sufficiently homogeneous populations. However it would seem wise to exercise caution and to ensure that ancestry effects are appropriately dealt with whenever the PRS or similar methods are employed. If the PRS did have some applicability, this would clearly be restricted to subjects having similar ancestry to that of the samples used to generate it. Given the results reported here one could not attempt to apply the PRS derived from European subjects to subjects with African ancestry. This highlights difficult questions about the fact that people of non-European ancestry may be disadvantaged if they cannot benefit from research which has been carried out largely or exclusively in European subjects. It is uncertain how much credence should be given to previous studies which may have used the PRS without appropriately accounting for ancestry effects. Although the current investigation has only dealt with the PRS for schizophrenia, it seems that similar considerations apply to the PRS for type 2 diabetes and coronary heart disease and this is likely to be the case for many other phenotypes (Reisberg *et al*., 2017).

## Acknowledgments

The author thanks the CommonMind, HapMap and PGC Consortia for making these datasets available. The CommonMind Consortium data generation was supported by funding from Takeda Pharmaceuticals Company Limited, F. Hoffman-La Roche Ltd and NIH grants R01MH085542, R01MH093725, P50MH066392, P50MH080405, R01MH097276, RO1-MH-075916, P50M096891, P50MH084053S1, R37MH057881 and R37MH057881S1, HHSN271201300031C, AG02219, AG05138 and MH06692. Brain tissue for the study was obtained from the following brain bank collections: the Mount Sinai NIH Brain and Tissue Repository, the University of Pennsylvania Alzheimers Disease Core Center, the University of Pittsburgh NeuroBioBank and Brain and Tissue Repositories and the NIMH Human Brain Collection Core. CMC Leadership: Pamela Sklar, Joseph Buxbaum (Icahn School of Medicine at Mount Sinai), Bernie Devlin, David Lewis (University of Pittsburgh), Raquel Gur, Chang-Gyu Hahn (University of Pennsylvania), Keisuke Hirai, Hiroyoshi Toyoshiba (Takeda Pharmaceuticals Company Limited), Enrico Domenici, Laurent Essioux (F. Hoffman-La Roche Ltd), Lara Mangravite, Mette Peters (Sage Bionetworks), Thomas Lehner, Barbara Lipska (NIMH).

## Ethics approval

This study was approved by the UCL Research Ethics Committee.

---

**Figure 2.**
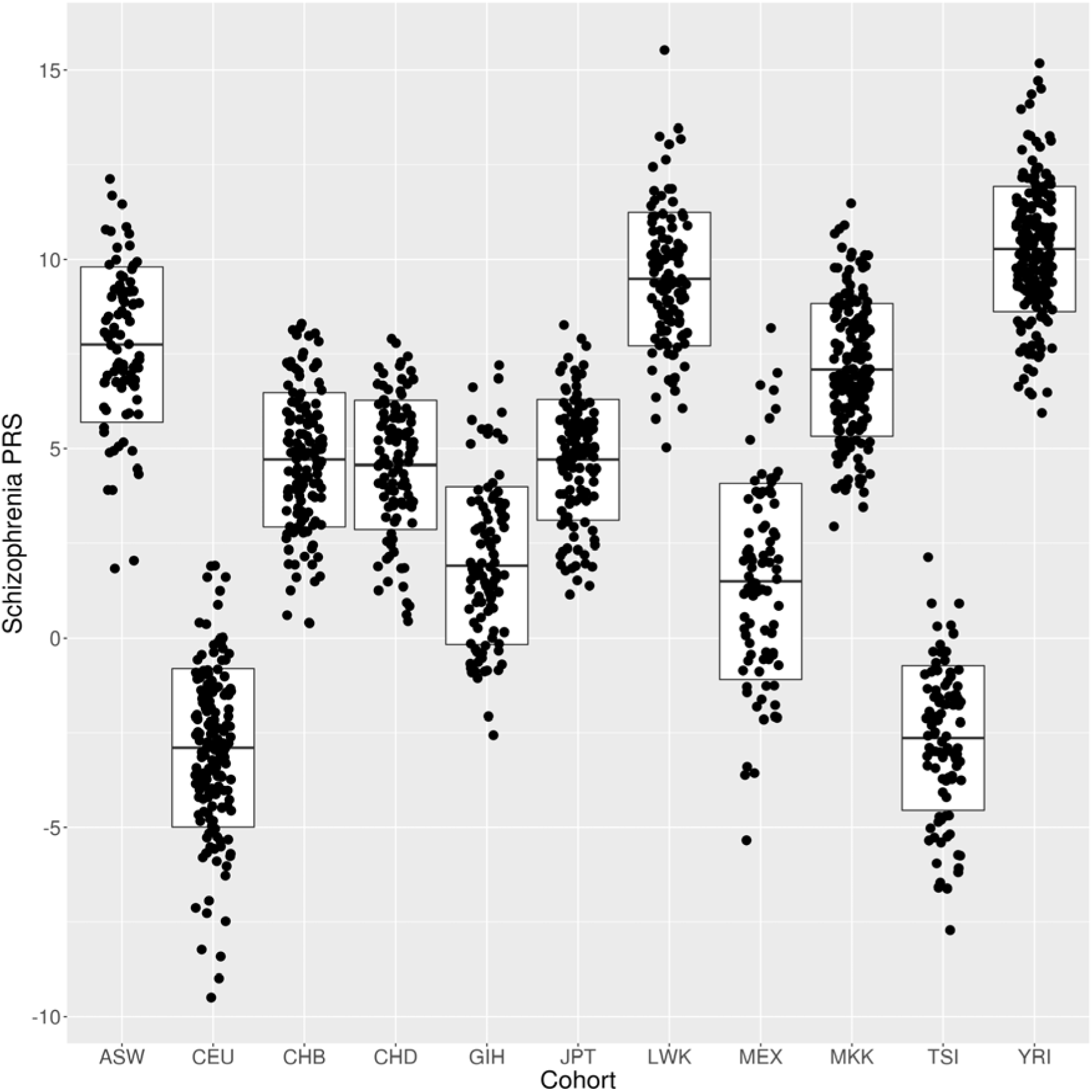
Schizophrenia PRS for individual HapMap subjects. The boxes show the mean and SD for each cohort.

